# Proteomics of highly post-translationally modified proteins in kidney tubulointerstitium

**DOI:** 10.64898/2026.09.30.755814

**Authors:** Jordan C. Langston, Alec Candib, Manveen Sethi, Joel Henderson, Joseph Zaia

**Affiliations:** Department of Biochemistry and Cell Biology, Boston University, Boston, MA, 02116, USA; Department of Pathology, Boston Medical Center, Boston, MA, 02116, USA

**Keywords:** Bioinformatics, Chronic kidney disease, Laser microdissection, Proteomics, Post-translational modifications

## Abstract

Kidney disease affects over 850 million people and leads to degradation of the kidney cortical compartments and kidney failure. Interstitial fibrosis and tubular atrophy (IFTA) is a universal feature of chronic kidney disease, yet the sequence of tubulointerstitial matrix protein (matrisome) changes that underlie the development and progression of IFTA are not fully understood. In particular, post-translational modifications which may stabilize key matrisome components and result in irreversible and functionally detrimental interstitial fibrosis or tubular basement membrane thickening, have yet to be characterized. Post-translational modifications (e.g., acetylation) have been shown to contribute to kidney diseases, yet they have not been investigated in chronic kidney disease progression. Herein, we established feasibility of matrisome protein quantification in the tubulointerstitium, with concurrent matrisome protein post-translational modification quantification, in archived diagnostic human kidney biopsies, using laser capture microdissection, liquid chromatography – tandem mass spectrometry (LC-MS/MS), and bioinformatics analysis. In this pilot study, we focused on two conditions associated with IFTA progression: diabetic nephropathy and kidney transplantation. We examined both healthy and chronically damaged tubulointerstitium in two human kidney biopsies – one representing each condition. We identified signature protein markers of tubulointerstitium (e.g., UMOD, SLC12A1) in abundance, confirming sample composition, and observed uniformity in protein abundance across replicates from the same biopsies. We found increases in certain collagen isoforms (COL1A1, COL6A3) and other ancillary proteins in areas of IFTA. We also successfully quantified post-translational modifications, confirming that hydroxylproline was the most abundant modification in healthy tubulointerstitium from the kidney transplant biopsy, and that hydroxyllysine was also highly abundant and could be confidently quantitated on two collagen isoforms (COL1A1, COL4A2). Lastly, we performed power analysis to determine the required sample size to achieve significant power and false discovery rate. Several proteins exhibited large Hedge effect sizes in the tubulointerstitium (ANPEP, PRODH2), indicating their potential usefulness as fibrosis signatures. Our study is a first step toward comprehensive characterization of tubulointerstitial matrisome changes, including post-translational modifications seen with IFTA progression. Subsequent studies will potentially reveal new molecular markers of key early events in chronic kidney disease progression, and new therapeutic targets for mitigating this process. Data are available via ProteomeXchange identifier PXD079025.

**Graphical Abstract:** 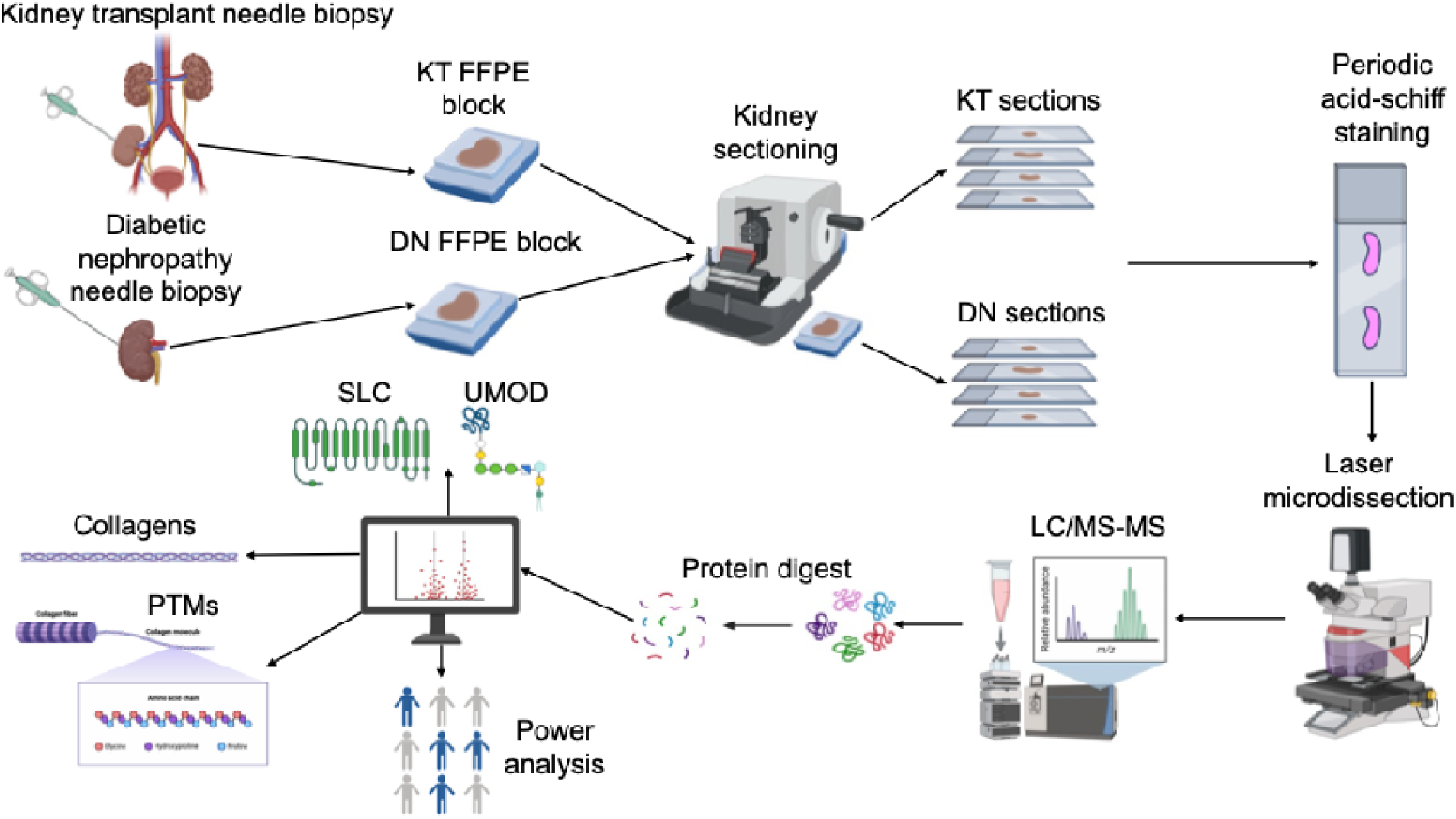

**Highlights:**

- Incorporation of spatial proteomics to study the tubulointerstitium
- Tubulointerstitium proteins exhibited the highest effect sizes in each comparison amongst kidney compartments.
- Greatest degree of hydroxyprolination amongst kidney transplant healthy samples compared to diabetic nephropathy.
- Computational analysis resulted in ∼50 samples in each biological group needed to obtain FDR < 0.05 and a power of 70%.

## Introduction

Kidney disease affects over 850 million people, equating to a global prevalence >10% (1). The number of people requiring dialysis and kidney transplantation is expected to increase from 2.6 million to 5.4 million in the next 5 years, costing over $85.4 billion annually in the U.S. alone(1). Kidney disease can, if left untreated, lead to an increased risk of other non-communicable iseases such as stroke, diabetes and cancer and chronic kidney disease (CKD)(2). CKD has a global prevalence of 11-13% population (3, 4), and it is projected to be the fifth leading cause of years of life lost globally (1). CKD results from irreversible damage to the kidney parenchyma, which includes the main kidney compartments (e.g., glomerulus, tubules and interstitium) in the renal cortex. In particular, the tubules and interstitium are two kidney compartments responsible for several kidney functions such as regulating blood pressure, immunity and erythropoiesis (5). Damage to the tubulointerstitium, as these two compartments are frequently combined into one, correlates with interstitial fibrosis and tubular atrophy (IFTA), which has been shown to be the final common pathway to end stage kidney disease (6), specifically in transplant patients and in diabetic nephropathy patients.

Transplant patients face elevated CKD risk due to suboptimal estimated glomerular filtration rate, allogenic factors including cell- or antibody-mediated rejection, or non-allogenic risk factors such as hypertension, diabetes bone mineral disorder and proteinuria. These factors may lead to endothelial cell dysfunction, arterial hyalinosis, glomerulosclerosis and eventual graft loss if not treated (7). Although systems-level canonical pathways of IFTA and CKD such as the renin-angiotensin-aldosterone system have been thoroughly explored, mechanisms that involve hypoxia, transforming growth factor β signaling (TGF-B), or cell senescence (8) remain poorly characterized. In particular, the elucidation of molecular mechanisms that alter the extracellular matrix proteome (hereafter the matrisome) in the kidney compartments, has not been investigated.

An increasing number of studies over the past 10 years have incorporated spatial biology methodologies (9), including those that use laser microdissection (LMD) with mass spectrometry (MS), to investigate the kidney (10–20) and kidney transplants (21). Examples of LMD studies focused on IgA nephropathy (22), diabetic kidney disease (DKD) (17), and chronic antibody mediated rejection (21) all of which contribute to CKD. Most of these studies focused on the glomeruli and identified differentially-expressed proteins. However, these studies did not investigate extracellular matrix (matrisome) pathology that can cause significant tissue parenchyma degeneration.

Matrisome dysregulation leads to fibrosis that is associated with 45% of deaths worldwide (23). The matrisome comprises an organized molecular network that modulates the filtration function of the kidney cortex (23). In particular, the matrisome in the tubulointerstitium and glomerulus consists of a 3-D network of core components such as fibrillar (e.g. types I, III) and network (e.g. type IV) collagens and glycoproteins (fibronectin, laminins, agrin, perlecan, versican) necessary for physiological parenchymal function (23). Collagens in the kidney provide the tensile strength for the basement membranes in each compartment (glomerulus, tubule, vasculature), support cell-matrix adhesion and are usually thickened during kidney disease, like other organs. **Supplementary Table 1** provides an overview of the collagens and their localization in each compartment based on a recent review (24). Agrin is the main heparan sulfate proteoglycan expressed in healthy kidneys, particularly in the glomeruli (4), and perlecan is increased in the glomeruli of IgA nephropathy patients (25). Elastin, another matrisome protein, is predominantly located in the vasculature (intima, media and adventitia) (3, 26). Sclerosis is the hardening or stiffening of the basement membrane because of fibrosis. In the kidney, the sclerosed glomerular basement membrane (GBM) and tubulointerstitium matrisome have increased collagen expression (e.g., types I, III, IV), and the hardened arterial GBM has increased proteoglycan (e.g., perlecan) expression in sclerosis (3). For example, over 1000 proteins have been annotated to the human matrisome (27) of which ∼140-200 are tissue-specific (28). Currently, 144 matrisome proteins have been identified in the glomerular matrix (10, 29), and 40-50 of these basement components localize in the GBM (29). In a subsequent study, proteomics of co-cultured GBM-associated podocytes and endothelial cells identified 127 matrisome proteins in endothelial cells and 142 in podocytes (30). Furthermore, in each study (10, 29, 30), network analysis was performed to visualize the interaction of different structural matrisome-associated and podocyte proteins, highlighting putative interactions and mechanisms contributing to disease. The tubulointerstitium matrisome has also been delineated, yet not to the same extent as the GBM. A set of 141 proteins have been identified in the tubulointerstitial matrisome (31), and although similar in composition to the GBM, the tubulointerstitial matrisome has a higher content of structural proteins such as vWF and EMILIN-2 (23). Thus, these findings defined the proteomes of the tubulointerstitial and GBM-associated podocytes and endothelial cells and suggested a role for cell–cell communication in the generation of the GBM and demonstrated that coculture of cells serves as a useful model for the *in vivo* GBM.

Recent review papers highlighted the roles of enzymatic and non-enzymatic PTMs of epigenetic modifiers in kidney disease, especially in DKD and IgA nephropathy (32, 33). Epigenetic PTMs acetylation and methylation exhibit complex effects on kidney diseases depending on the context. For example, acetylation correlates with inflammation in CKD, glomerular fibrosis in DKD (32, 34) and reduced matrisome protein expression in CKD (35). Glycosylation motifs can also contribute to kidney pathophysiology (36). For example, N-glycosylation of ENTPD5 promoted apoptosis of kidney tubule epithelial cells in DKD (37). However, non-epigenetic PTMs have also shown to contribute to tissue physiology, and these targets, especially hydroxyproline, are highly abundant in diseased vs healthy samples. We employed peptide-level analysis to investigate changes in abundances between biological groups, potentially indicating disease-related mechanisms. Furthermore, the analysis of PTMs can reveal altered site localizations and differences in abundances between groups. PTMs are crucial biochemical modifications that could affect the downstream function of a cell’s proteoform. Over 400 different PTMs exist, thus regulating a myriad of diverse physiological (e.g., cell adhesion/migration) and pathophysiological (e.g., inflammation) cellular processes and regulatory mechanisms (38). Most PTMs are reversible and added enzymatically to amino acids, yet others are regulated non-enzymatically (34). Furthermore, PTM modification can lead to the formation of structural isomers; for example, hydroxylation of proline can occur on the 3- or 4 position (39) and thus contribute to unique biological mechanisms, including in the kidney (40). However, most (LMD) proteomics studies fail to investigate and report on PTM identification on the protein or peptide level. PTMs contain a wealth of information necessary to comprehensively characterize a cell’s proteoform using advanced bioinformatics methods for understanding protein structure and function.

Hydroxylation occurs to 10-13% of collagen proline residues and is essential for the structural integrity, thermal stability, and mechanical function of the triple helix. Collagens are a main component of the increased matrisome that leads to fibrosis (41) and hydroxylproline secreted in urinary microvesicles is a marker for kidney fibrosis in transplant recipients (40). There is a compelling need to consider the roles of collagen hydroxylproline levels and fibrosis in the kidney compartments (e.g., tubulointerstitium) in disease progression. However, there remains a significant dearth of knowledge in assigning PTMs to collagens and other matrisome proteins in kidney disease(s) to determine pathological alterations that contribute to CKD. Thus, in this study, we not only perform LMD-MS proteomics on the tubulointerstitium from cases two common kidney diseases (transplant and diabetic nephropathy) but also characterize and quantify PTMs on the protein and peptide level in the matrisome using advanced bioinformatics methods.

## Materials and Methods

### Study design and human kidney tissue preparation

De-identified formalin-fixed paraffin embedded (FFPE) blocks of archived diagnostic human kidney biopsy material were retrieved retrospectively from the Boston Medical Center (BMC) Department of Pathology and Laboratory Medicine. This study of deidentified human biospecimens was exempted from IRB review (H-45508). Tissue sections prepared from this material were evaluated to identify appropriate pilot specimens for this study. We sought to ensure sufficient cortical tissue was present for laser capture microdissection, protein isolation and quantification, and that regions representing both healthy tubulointerstitium (i.e. no chronic damage present), and areas of chronic damage (i.e. IFTA), were present in sufficient quantities in each biopsy utilized for this pilot study.

We chose two de-identified kidney biopsy cases for this pilot study: a native kidney biopsy exhibiting features of moderate diabetic nephropathy (DN), and a kidney transplant (KTRH) case with no evidence of rejection and minimal chronic damage. Thus, for this pilot study, N=1 for each biological group. The DN case was specifically chosen to not exhibit any underlying pathologic processes (e.g., acute tubular injury) in the tubulointerstitium that would confound results. Similarly, the kidney transplant case was devoid of any underlying pathologic processes affecting the tubulointerstitium (e.g., rejection, inflammatory or infectious mechanisms, etc.) and was chosen to serve as a “positive control” for comparison to both healthy and chronically-damaged parenchyma in the DN sample. From these two kidney biopsies, tissue representing three distinct states or “biological groups” of tubulointerstitium were examined: healthy tubulointerstitium from the DN biopsy (“DNH” - diabetic nephropathy healthy); chronically damaged tubulointerstitium from the DN biopsy (“DNS” - diabetic nephropathy severe); and healthy tubulointerstitium from the transplant kidney biopsy (“KTRH” - kidney transplant healthy). **Table 1** illustrates demographics for the two coded and deidentified cases.

**Table 1.**
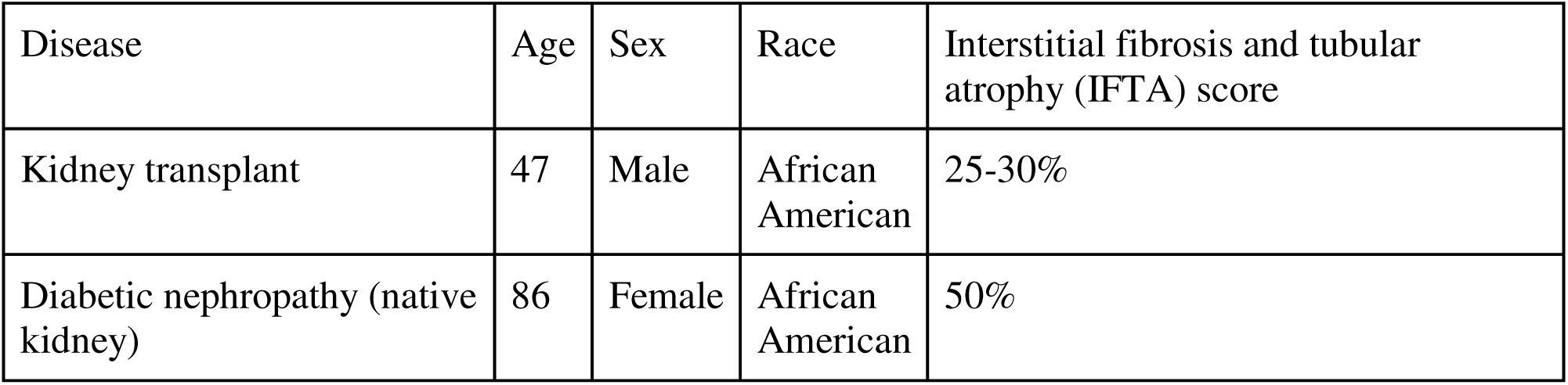
Demographics of the two cases used in this study. General characteristics of the patients are shown. Interstitial fibrosis and tubular atrophy scores were obtained from BMC’s Department of Pathology and Laboratory Medicine database.

| Disease | Age | Sex | Race | Interstitial fibrosis and tubular atrophy (IFTA) score |
| --- | --- | --- | --- | --- |
| Kidney transplant | 47 | Male | African American | 25-30% |
| Diabetic nephropathy (native kidney) | 86 | Female | African American | 50% |

### Sectioning of FFPE blocks

We used a Leica RM-2235 microtome for sectioning of FFPE blocks. The thickness of each section was set to 4 μm to enhance the histological identification of the tubulointerstitium. A set of serial sections from each FFPE block represented a single replicate. Once sectioned, ribbons were placed in a Tanner scientific 450 water bath (New England Biomedical Services) set at 44°C to dissociate individual sections. Tissue sections were mounted on polyethylene napthalate (PEN) coated slides (Leica, 2 µm, No. 11505158, batch 000703-25). A maximum of three sections were placed onto each slide to prevent the dissociation of sections from the PEN membrane. In some cases, serial sections from a block were adhered to different PEN slides. For example, if 6 serial sections were cut from a block and 3 sections were placed on one slide and 3 on another, then both slides would be considered the same technical replicate. To ensure we had adequate technical replicate for LMD, a minimum of 6 serial sections was defined as a technical replicate. To facilitate assessment of intra-group variability, three to four technical replicates were prepared for each of the three tubulointerstitial biological groups (DNH, DNS, KTRH).

### Staining and imaging of PEN slides

Once sections were adhered to the PEN membrane, staining of the slides was performed using the Abcam Periodic-acid Schiff (PAS) staining kit (Fisher Scientific, No. NC1642221). FFPE slides were deparaffinized according to the following protocol (42). Slides were submerged twice in xylenes (Fisherbrand HistoPrep Xylene, Lot 152513) for 3 minutes for dewaxing. Graded alcohol solutions (Decon Laboratories, Ethanol 200 PROOF, CAS 64-17-5, Lot# 2772314) and distilled water were implemented for tissue rehydration. For rehydration of the tissue slides, slides were submerged in these successive steps: twice in 100% ethanol for 1 minute, 95% ethanol for 1 minute, 70% ethanol for 1 minute, 100% distilled water for 1 minute. Slides were immersed in Periodic-acid Schiff (PAS) for 10 minutes, and then briefly rinsed in distilled water. Slides were then placed in Schiff’s solution for 30 minutes. Lastly, slides were dehydrated by submerging the slides in 95% ethanol for a minute and then 100% ethanol for another minute. Samples were placed in the -80°C until LMD. Whole slide images were generated at 40x using Leica Laser Application Suite X (LAS-X) software.

### Laser microdissection

We used the Leica LMD 6 system for performing laser microdissection of tissue sections from each technical replicate. Prior to LMD, samples were left to dry in a fume hood for one hour to facilitate LMD. We microdissected 2.5 million µm³ (or ∼625,000 μm^2^) of 4 μm thick tissue slices of tubulointerstitium from each technical replicate into 0.5 mL PCR tubes. These tubes were pre-washed with methanol, ethanol and LC-MS water to minimize MS interference. A 60 µL volume of Optima liquid chromatography (LC)-mass spectrometry (MS) water from Fisher Scientific (Waltham, MA, Lot# 245317) was added to the cap prior to tissue collection. We used a magnification of 63x for LMD to distinguish between healthy-appearing and diseased-appearing tubulointerstitium morphology. Healthy appearing tubulointerstitium adhered to the following criteria: presence of a brush border in the tubular columnar epithelial cells, relatively free of cell debris, minimal interstitial expansion, absence of inflammatory cells in the tubules and thin tubular basement membranes. Chronically damaged tubulointerstitium had severe tubular atrophy, thick tubular basement membranes, extracellular matrix (e.g., collagen) deposition, presence of inflammatory cells in the tubules and/or interstitium, tubular epithelial cell flattening. Additionally, we consulted with a renal pathologist to confirm tubulointerstitium structural features. Regions of interest (ROIs) for LMD were drawn free-hand and each ROI had an average area ∼8000 μm^2^ (32,000 µm³). Tubules and interstitium were collected in a single tube for each replicate. Laser power for LMD ranged from 20-30 and an average speed of 30 was incorporated to achieve a clean extraction of each ROI. After collection, we stored samples at - 80°C prior to digestion. We captured whole slide images of the two cases (**Figure 1**). Only healthy regions were collected in the KTRH case, while healthy and chronically damaged regions were collected from the DN case.

**Figure 1:**
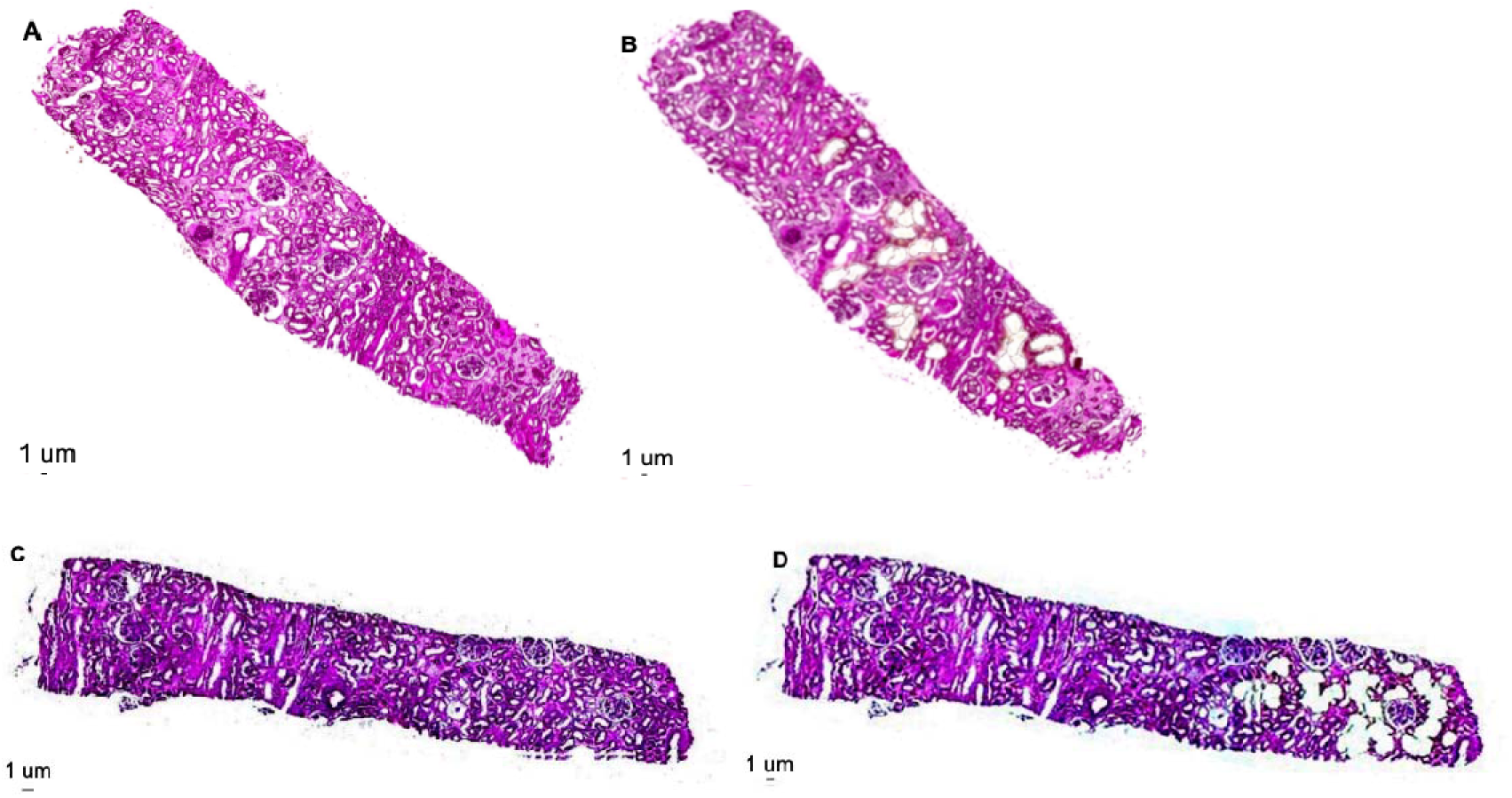
Whole slide images of cases. Examples of whole slide images of the DN kidney case before LMD (Panel A) and after LMD (Panel B), and a KTRH case before (panel C) and after (panel D) LMD. In the DN case, healthy and chronically damaged regions are represented, yet the areas collected were distributed to two separate replicates. Cases were stained with periodic acid Schiff to visualize compartments and imaged at 40x. Respective scale bars are shown.

### Protein digestion

Ammonium bicarbonate (ABC) (Sigma Aldrich ReagentPlus ≥ 99.0%, Lot# BCBV3862, A6141-KG), trypsin/lysine-cysteine (Lys-C) (Promega Mass Spec Grade Trypsin/Lys-C, Fisher Scientific, Catalog No. PRV5072), chondroitinase ABC (Chondroitinase ABC from *Proteus vulgaris*, Sigma-Aldrich), trifluoroacetic acid (TFA) (FisherChemical, Lot 230768, A116-10X1AMP, OptimaLC-MS, CAS 76-05-1), enolase from baker’s yeast (*S. cerevisiae*, E6126-500UN, Sigma Aldrich, Catalog No. 50-178-0627) and acetonitrile (FisherChemical, Lot# 245936, A955-1, OptimaLC-MS, CAS-75-05-8) were purchased for protein and glycosaminoglycan (GAG) digestion of the LMD tubulointerstitium. We prepared a 100 mM solution of ABC in deionized water and a 1 µg/µL stock solution of enzyme Lys-C and trypsin. After samples were removed from the freezer, we added ABC to each sample, vortexed and centrifuged (Brinkmann Instruments Eppendorf Centrifuge 5451D) samples at 2,000 g for 10 minutes. We heated samples for an hour at 95°C on a thermomixer (Eppendorf Thermomixer C Model 5382) to initiate denaturation, followed by another hour at 75°C in 12% v/v of ACN. We removed samples from the thermomixer, allowed them to cool for 15 minutes prior to the addition of 2 mU chondroitinase ABC in a 100 mM ammonium acetate buffer. Chondroitinase ABC was used to cleave chondroitin GAG chains. We placed samples on the thermomixer at 37 °C for two hours. We removed samples from the thermomixer and added 16 ng of yeast enolase per μg of sample protein, to capture sample processing variation from tryptic digestion and LC-MS. We added 100 ng/µL of trypsin and Lysine-Cys to each sample, encapsulated the samples tubes in parafilm to prevent evaporation and placed them onto the thermomixer overnight (37°C). The next day, we removed samples from the thermomixer and added 1% v/v of TFA to stop digestion. We placed samples in a speedvac (Thermosavant SPD1010 SpeedVac system) to dried and subsequently stored them in a - 20°C freezer prior to cleaning.

### Peptide cleaning

We incorporated Thermo Scientific Pierce C-18 ziptips (Lot AB401129, Ref 89870) for cleaning and desalting peptides in each sample. We prepared for peptide cleaning: 50% ACN/50% water, 2% ACN/0.1% TFA and 60% ACN/0.1% TFA. Before cleaning, ziptips were tapped on the bench to activate the resin. We originally placed ziptips in the wash tubes for each sample. We pipetted 50% ACN/50% into each column, centrifuged at 1,000g for 1 minute. The wash was discarded. We injected 2% ACN/0.1% TFA to each column, centrifuged and repeated two more times. Each wash was discarded. We transferred columns to the flowthrough tubes. We removed the peptides from the -20 C freezer and resuspended them in 2% ACN/0.1% TFA. We vortexed peptides (VWR Mini Vortexer Scientific Products), centrifuged and applied them to the equilibrated columns. We added 2% ACN/0.1% TFA to the sample peptide tubes, vortexed, centrifuged and applied them to columns. After centrifugation, the flowthrough was applied, centrifuged, reapplied (twice) and centrifuged again. We transferred columns to the wash tubes, and each column was washed with 60 µL 2% ACN/0.1% TFA. Each wash was discarded. Columns were transferred to peptide tubes and washed via centrifugation thrice with 30 µL 60% ACN/1% TFA. All washes were collected in the same tube. Clean peptide tubes and flowthrough tubes were dried down and stored at - 20°C prior to LC-MS analysis.

### Liquid chromatography-tandem mass spectrometry analysis

Peptides were removed from the -20°C. 5 μL of mobile phase A (99% water/1% acetonitrile/0.1% formic acid) was added to each sample. Samples were then centrifuged for 10 mins at 10,000 g at 4°C. Pierce retention time calibration mixture (43) and Hela cell lysate digests were added before injection of samples as external quality controls. The peptides were loaded onto a Thermo Scientific™ Orbitrap Fusion™ Lumos™ Tribrid™ mass spectrometer (MS). Nano-LC-MS/MS separation was performed using a NanoAcquity M-Class UPLC (Waters Technology) coupled to an Orbitrap Fusion Lumos MS (Thermo Fisher Scientific). Reversed-phase nanoEase V/M Symmetry C18 trap column with a 180 μm internal diameter and a nanoEase M/Z HSS T3 column with a 75 μm internal diameter were used with a 75 min method at a constant flow rate of 0.5 µL/min using mobile phase A and mobile phase B (99% acetonitrile/1% water/0.1% formic acid). The data dependent acquisition (DDA) was performed in positive mode over a mass range of m/z 350-2000, with a maximum injection time of 50 ms and a resolution of 120 K. The normalized automatic gain control (AGC) target was set to 100% (4e^5^) and dynamic exclusion to 10s. Tandem MS was performed using an isolation window of m/z 1.6, a microscan of 1, a resolution of 30 K, a maximum injection time of 60 ms, and an AGC target of 5e^5^ at 1000%. A fixed normalized collision energy (NCE) in the high collision energy dissociation (HCD) mode of 27 V was used. All samples were handled, processed and analyzed simultaneously to avoid instrument bias and batch effects. Furthermore, samples were randomized to control for instrument variability.

### PEAKS bioinformatics analysis

We used PEAKS Studio 13.1 (Bioinformatics Solutions Inc, Waterloo, ON, Canada) for protein, peptide and post-translational modification (PTM) identification and quantification. Technical replicates were grouped as follows prior to label-free quantification (LFQ) analysis: diabetic nephropathy severe (DNS), diabetic nephropathy healthy (DNH) and kidney transplant recipient healthy (KTRH). Trypsin was chosen for *in silico* digestion, data-dependent acquisition (DDA) was the acquisition method, high energy collisional dissociation (HCD) was the fragmentation pattern for each sample. The “proteome (*de novo* assisted proteomic analysis)” was the chosen workflow. Data refine options were set to default. Precursor mass was 10 ppm and fragment ion 0.02 Da. Enzyme was set to trypsin, digest mode was semi-specific and missed cleavage was set to 2. Carbamidomethylation and oxidation of methionine were fixed PTMs. Variable PTMs included hydroxylproline, hydroxylysine, hydroxylysine-hexose, hydroxylysine-hexose-hexose. Taxonomy was set to Homo sapiens and Saccharomyces. The target database included a XML file of all the protein coding genes from Uniprot and a contaminant database containing a list of all known contaminants from Uniprot (release 2026_01). Peptide length was set from 6 to 45. All other parameters were set to default. Four PTMs per peptide were permitted for variable search. Quality control was enabled. Samples were normalized using yeast enolase 1 (ENO1). “Global” normalization strategy was incorporated. The number of unique peptides per protein was >= 2. Regarding data filtering for LFQ quantification, the following parameters were used: significance 15, fold change 1-64, feature intensity >=0, average area 1E9, charges from 1-10 and the top 3 peptides were used for protein quantitation. To ensure confident PTM localization analysis with 99% certainty, we retained those PTMs that had an ambiguity score (or AScore) >= 20, correlating with a p < 0.01 (44).

### RStudio analysis and visualization

We imported the normalized label-free quantified (LFQ) protein and protein-peptide files into RStudio (41) (4.5.2) for statistical and bioinformatics analysis. We removed any contaminants from the protein list. To ensure reproducibility of protein and peptide quantitation, we kept proteins and peptides that had quantitation in 70% of technical replicates within one biological group for downstream analysis (45). Datasets were normalized via log2 transformation and proteins with missing values were imputed with a value one standard deviation below the minimum intensity within each sample to mimic MS background. Post-translational modifications (e.g., hydroxproline, hydroxylysine) were quantified as well, and we only used those modifications that had an AScore >= 20, corresponding to a p-value < 0.01 (44). This filtered dataset was used for all downstream analyses. We used the stats and corrplot packages to perform hierarchical clustering and correlation analysis and ggplot2 for data visualization. We incorporated ComplexHeatmap packages to generate heatmaps of the log2 normalized abundances of proteins and peptides. **Supplementary Excel File 1** contains the list of quantified normalized proteins and peptides (before and after imputation). **Supplementary Excel File 2** contains the list of PTMs identified and quantified across samples on protein- and peptide levels.

### Quantitation of kidney compartmental-specific proteins

To determine the content of tubulointerstitium and glomerulus in each microdissected sample, we identified compartment-specific proteins using the Kidney Precision Medicine Project (KPMP) database (46) (47). KPMP is an ongoing, NIH funded, multi-consortium study elucidating the underlying molecular mechanisms in kidney disease. We investigated KPMP’s LMD proteomics data to identify and confirm significantly enriched tubulointerstitium and glomerular proteins in our data as those possessing an adjusted p < 0.05 and fold change > 0 when comparing compartments. Results from our study were compared to KPMP’s published atlas for validation. We also used Human Protein Atlas (48) and literature to investigate the functions of these proteins and confirm those that were true signatures in each compartment.

### Power analysis

Due to N=1 of a sample within a biological group (DN and KTRH) and the incorporation of technical replicates, standard statistical tests (e.g., t-tests) were not used for inferential analysis. Results from statistical analysis would not represent true biological variation across the population. Thus, we calculated descriptive statistics (log2 fold changes and Hedge’s g effect sizes) for each quantified protein between comparisons. We chose Hedge’s effect size and not Cohen’s due to the small sample size, and Hedge’s applies a bias-correction factor to mitigate inflation that occurs in small-scale studies. An effect size (g) > 0.2 was considered small, indicating groups appeared the same. A g > 0.5 correlated with a moderate indicative of noticeable differences, while a g > 0.8 inferred a large effect and clear separation between groups (49). The incorporation of effect sizes in proteomics datasets as an additional, statistical metric to evaluate data other than inferential p-values is emerging in new proteomics packages (50). Hedge’s g represents the effect size of each protein amongst comparisons. It describes the standardized change in magnitude a feature or variable experiences between two groups (e.g., control vs treatment). It is calculated as follow:

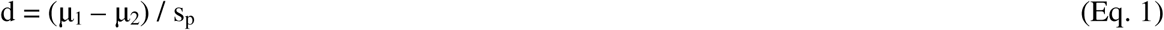

where μ_1_and μ_2_ represent the group means of a protein’s expression, s_p_ is the pooled standard deviation and d is Cohen’s d. This value is then multiplied by a correction factor (J) to adjust for small size bias and prevent overestimation of sample size:

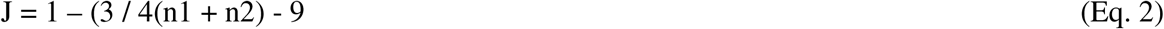

to produce the final equation

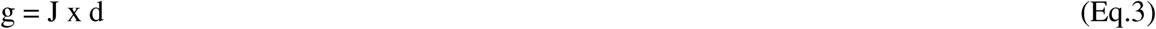

where g is Hedge’s effect size, and n is sample size.

We created a limma-based model (51) in RStudio to determine the number of biospecimens in each biological group necessary to obtain a FDR < 0.05% and a power of 70% per comparison. Limma calculates differential expression proteins by using an empirical Bayes model to shrink gene/protein variance to a constant value that stabilizes protein expression across samples. We chose 70% power and not the conventional 80% due to the preliminary nature of this study. The model incorporated the list of effect sizes and means calculated for each protein in each comparison and the raw protein-specific variability/residuals from the regression predictions. We assumed 10% of proteins would be differential expressed in a future study. Due to the incorporation of technical replicates in our study, variance was underestimated and artificially low. We used variance inflation factors ranged from 1-4 for each comparison. We expected to observe a decrease in power as variance increases, thus requiring more samples to observe a true biological change in the population. Our bootstrapped model simulated 1000 iterations using these parameters to compute FDR and power for each comparison.

## Results

### Proteomics analysis highlights tubulointerstitium reproducibility within biological groups

Liquid chromatography-tandem mass spectrometry (LC-MS/MS) was used to analyze the samples (hereafter technical replicates) resulting in 11 LC-MS/MS runs. Total ion chromatograms for each sample, are provided in **Supplementary Figure 1**. We incorporated internal, external and system standards to correct for variability in systems performance and for comparison of protein abundances between technical replicates (43). Technical replicates were grouped as follow: Group 1: Diabetic nephropathy severe (DNS); Group 2: Diabetic nephropathy healthy (DNH) and Group 3: Kidney transplant healthy (KTRH). Prior to any downstream analysis, we first evaluated yeast ENO1 reproducibility across MS runs. Since we added a consistent ratio of yeast ENO1 to microdissected tissue volume for each sample, we expected variability to be low, thus permitting normalization of proteins. The coefficient of variation (CV) was 0.3 for yeast ENO1. Furthermore, proteins with increasing log-concentration tended to have lower CV, narrow spread of CVs and adhere to a linear pattern (**Figure 2**) (52).

**Figure 2:**
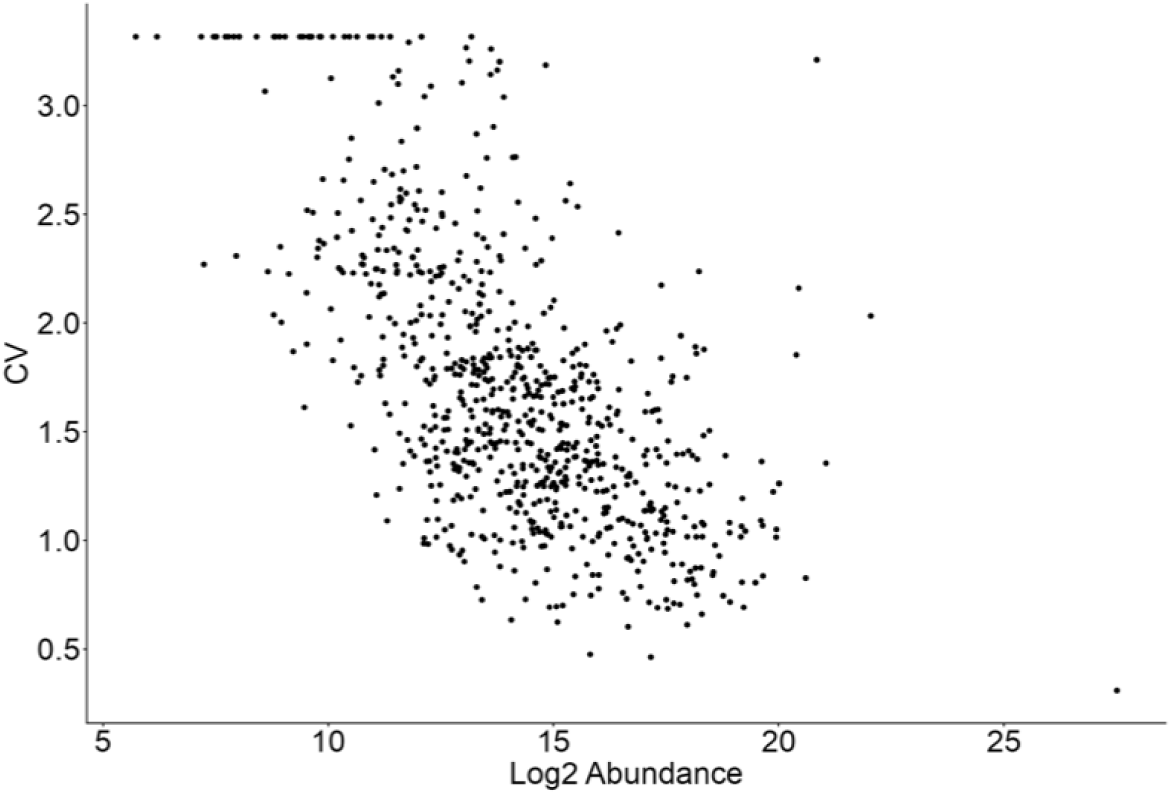
Protein abundance variability across technical replicates. Scatterplot modeling changes in log2 normalized abundance as a function of coefficient of variation.

After normalizing protein abundance to yeast ENO1, we performed quality control analysis to determine if unique clustering patterns were present (Figure 3).

**Figure 3:**
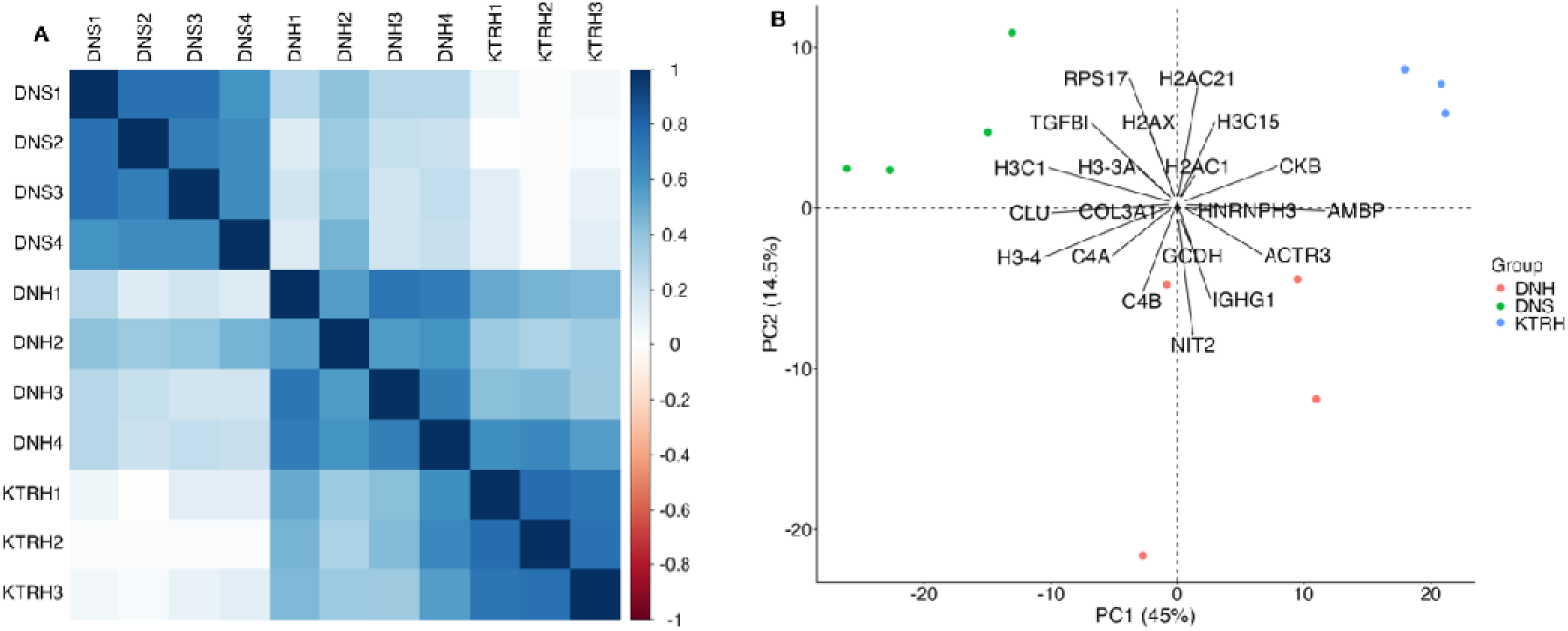
Clustering of kidney tissue technical replicates. Panel A illustrates a correlogram highlighting Pearson correlation coefficients between technical replicates, with blue highlighting a correlation = 1 and red highlighting a correlation = -1. Panel B shows principal component analysis of the technical replicates. Green dots = Diabetic nephropathy severe (DNS). Blue dots = Kidney transplant healthy (KTRH). Red dots = Diabetic nephropathy healthy (DNH).

Correlogram analysis (**Figure 3A**) highlighted uniform correlation within a biological group and low correlation between groups, demonstrating high quality and consistent protein expression within groups. Principal component analysis (PCA) (**Figure 3B**) illustrated separation of technical replicates within each biological group with PC1 exhibiting greater separation power than PC2. Additionally, the top 20 proteins driving separation in the PC1-PC2 plane were also shown. Four of these top drivers (alpha-1-microglobulin/bikunin precursor (AMBP), clusterin (CLU), collagen type 3-alpha 1 chain (COL3A1), transforming growth factor beta induced protein (TGFBI)) were tubulointerstitium proteins (31). CLU, COL3A1 and TGFBI exhibited greater association in the DNS replicates, while AMBP illustrated higher association in the KTRH replicates. Thus, these QC metrics indicated that LMD regions from each case were captured with high fidelity and reproducibility precluding cross-interference during MS. Confirmation of normalized proteins via yeast ENO1 and intra-group homogeneity allowed us to confidently identify and quantify proteins in our pilot study. We identified 852 protein groups corresponding to 817 unique proteins and 6516 peptides with ≥ 2 unique peptides/protein. Furthermore, using our criteria to ensure reproducibility in protein quantitation, we included 671 quantified proteins and 2821 peptides for downstream analysis. We were then interested in determining whether these proteins were signature or non-signature tubulointerstitium proteins, and if there were non-tubulointerstitium (glomeruli or “other”) proteins. Signature proteins are known cell expression markers characterizing unique cellular functionality, while non-signature proteins are typical housekeeping proteins yet have been shown to play an important role in a cell. Although LMD is a highly reproducible technique, it is possible that non-tubulointerstitium compartments were captured due to the relative proximity of each compartment with another. Thus, we investigated compartmental-specific proteins in our pilot data.

### Quantitation of compartmental-enriched proteins highlights signature and non-signature proteins

We estimated the percentage of each compartment (e.g., glomeruli, tubulointerstitium and non-compartmental enriched (“other”) proteins) that was captured for each sample replicate in our normalized quantified protein list. As stated, we first identified compartmental significantly enriched proteins in the KPMP database based on our criteria and pinpointed which ones were detected in our data. Once compartmental proteins were confirmed, we calculated the ratio of the number of compartment proteins identified in each sample by the total number of quantified proteins in each sample. Since we only captured tubulointerstitium (as a combined compartment) in this pilot study, we expected to identify and quantify more proteins associated with tubulointerstitium compared to those specifically associated with glomeruli. We also expected to identify an abundance of non-specific proteins that were not enriched in a compartment, termed “other”. **Figure 3** illustrates the identification and abundances of compartmental-enriched (signature and non-signature) and non-specific proteins.

**Figure 3:**
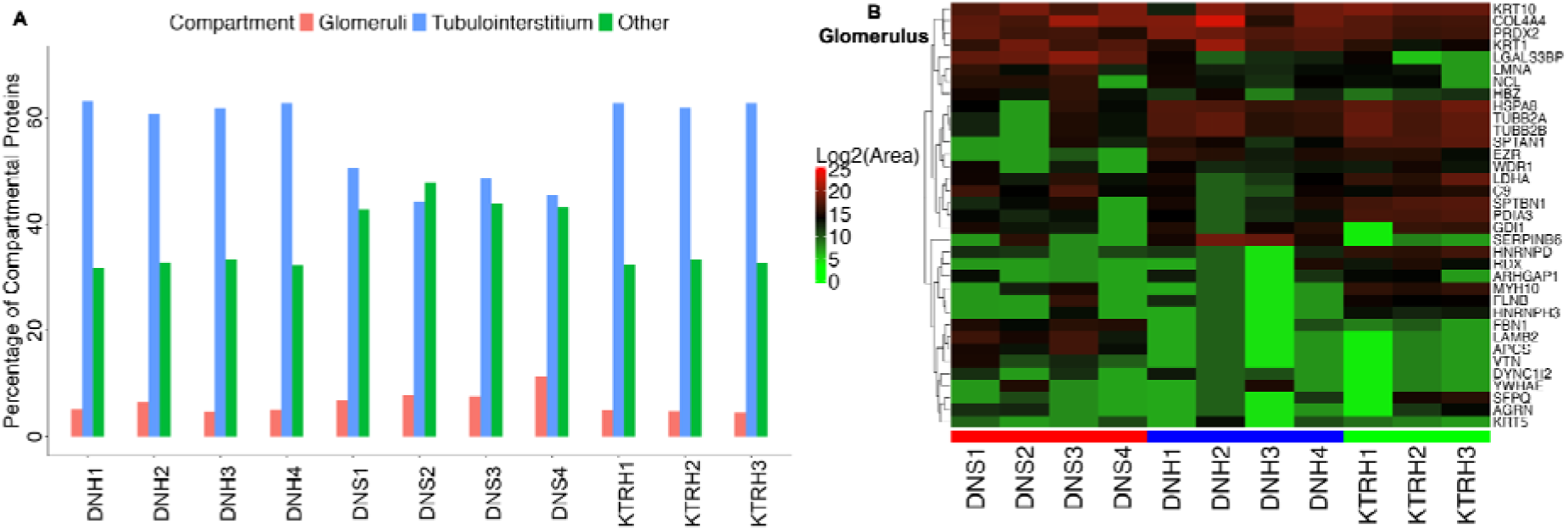
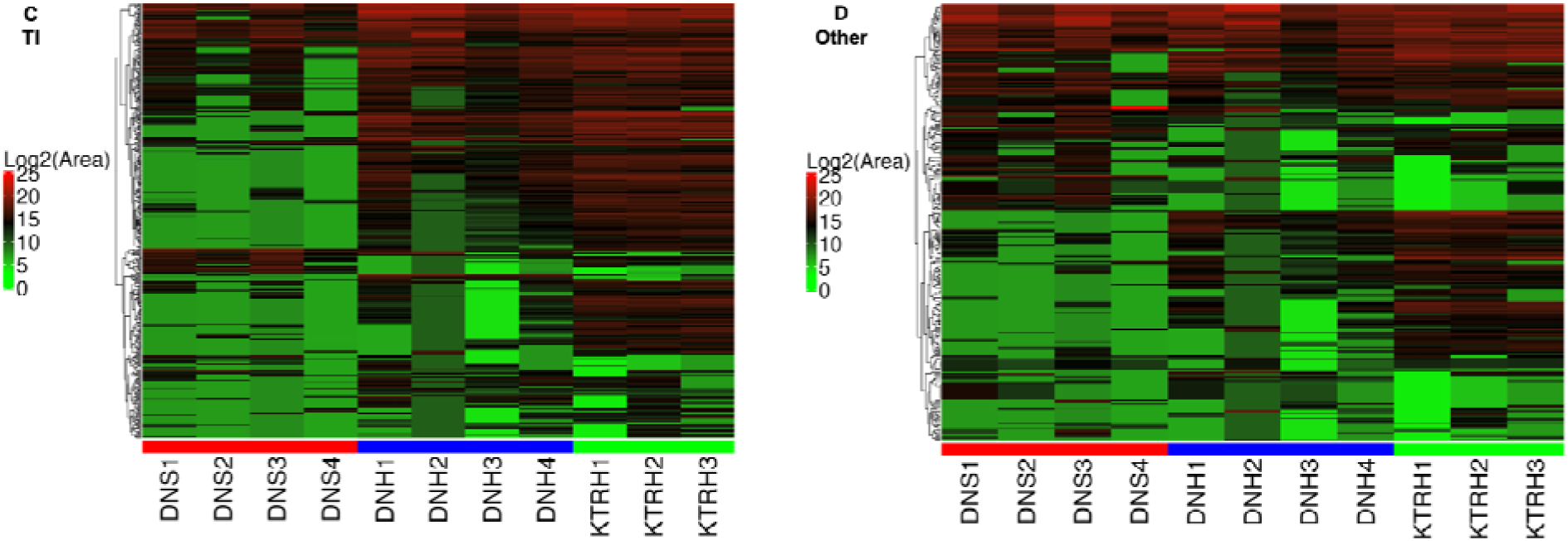
Quantitation of compartmental-specific proteins. Proteins associated with the glomeruli, tubulointerstitium and “other” proteins were identified across technical replicates representing each of the three biological groups. Panel A highlights the percentage of compartmental-specific proteins in each sample. Panels B-D illustrate the unsupervised hierarchical clustering of summed log2 scaled abundances of glomerulus, tubulointerstitium (TI), and “other” proteins, respectively, across replicates. In panel A, red bars = glomerular proteins, purple bars = tubulointerstitium proteins, and green bars = “other” proteins. In panels B-E, green bars = DNS samples, red bars = DNH samples and blue bars = KTRH samples. The color key on the left in each panel represents the normalized log2 scaled abundance. Due to the multitude of “other” and tubulointerstitium proteins, proteins are listed in **Supplementary Excel File 3**.

These results showed that the technical replicates did not include significant quantities of glomerular proteins. As shown in panel A, most proteins in our quantitated proteomics data were categorized as tubulointerstitium or “other”, indicative of the consistency in LMD across all replicates regardless of disease state. KTRH replicates had the highest percentage of tubulointerstitium proteins identified, while DNS replicates had the lowest. There were no substantial differences between groups for glomerular and “other” proteins, yet differences were observed for the tubulointerstitium proteins. “Other” proteins in our proteomics data possess potential housekeeping roles, or they could be present in more than one compartment, but they were not significantly enriched. Thus, they were excluded as we focused on compartment enriched proteins.

Signature proteins were not identified in the glomerulus, but they were identified in the tubulointerstitium such as solute carrier family transporters (SLC12A1), megalin (LRP2) and cubilin (CUBN) (53). SLC12A1 was highly abundant in the DNS group compared to the DNH and KTRH replicates. Tamm-Horsfall or uromodulin (UMOD) protein is another classical tubulointerstitium protein (54, 55). UMOD was more abundant across all DNS replicates compared to DNH and KTRH replicates. We also observed protein abundances associated with IFTA (DNS) compared to non-IFTA regions. For example, COL6A3 was more abundant across all DNS replicates; furthermore, COL14A1 and COL5A1 exhibited minimal abundance across most replicates except for two DNS replicates. Tenascin-C (TNC) was more abundant across DNS replicates compared to KTRH and DNH replicates. Fibrillar collagens (COL1A1, COL1A2, COL5A1) and non-fibrillar collagens (COL6A3, COL14A1). COL1A1 and COL1A2 exhibited uniform abundance across most replicates highlighting the presence of excess collagen deposition in these replicates and matrisome turnover dysfunction. It is important to note that our quantified data in this pilot study, due to small sample size, only contained a subset of the total number of unique enriched proteins associated with a compartment. Due to the abundance of collagens in our pilot data across all biological groups, we quantified post-translational modifications on the protein-level to determine if there were substantial alterations.

### Protein-level post-translational modification analysis highlights hydroxyproline and hydroxylysine as highly abundant

We defined the percentage of confident post-translational modifications (PTMs) on the protein level by taking the ratio of confident modification sites by the total number of modification sites and examining distribution of PTMs across replicates as shown in **Figure 4**. As shown, hydroxyproline (HPro) had the highest ratio of confident PTM sites to total PTM sites, followed by hydroxylysine (HLys). The abundances of N-acetyl-hexosamine-hexosamine (HexNAcHex) and N-acetyl-hexosamine (HexNAc) modifications, indicating glycosylation, was lowest in each technical replicate, and analysis of MS did not indicate any potential fragmentation moieties associated with significant glycosylation (**Figure 4A**). Across all replicates, HPro and HLys protein abundance is higher in KTRH and DNH replicates compared to DNS (**Figure 4B**). The investigation of PTMs on a protein level only represent an overview of its underlying biology. Molecular alterations on the peptide level would yield greater insight on how PTM abundance is shifting during disease, leading to new mechanistic studies evaluating their (patho)physiological roles.

**Figure 4:**
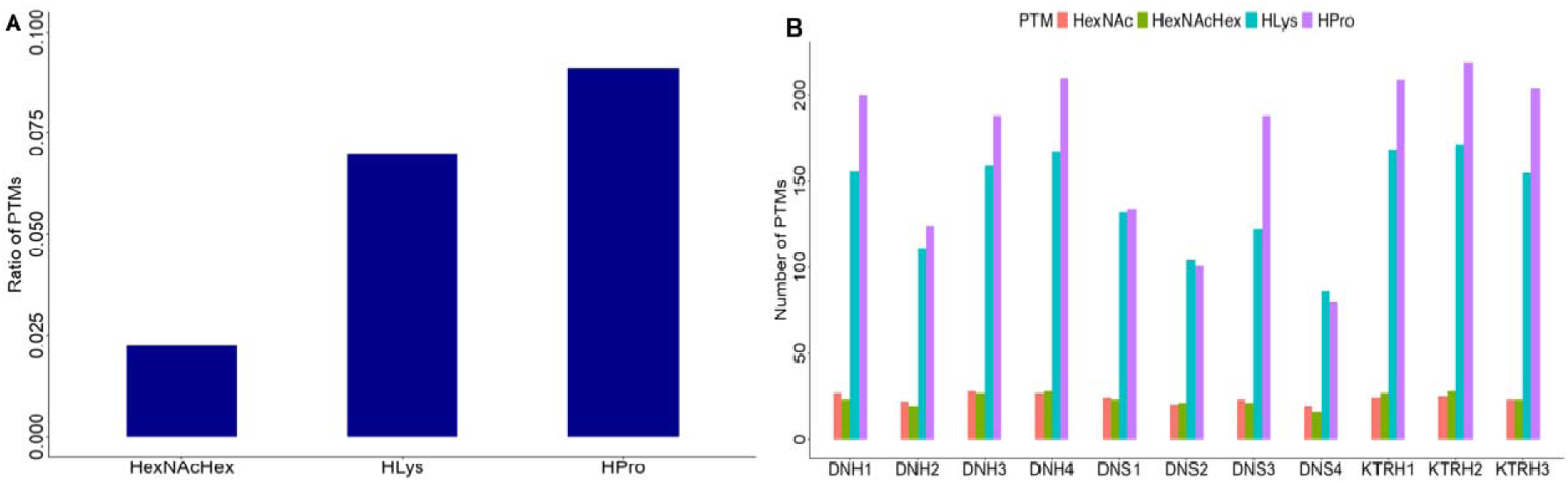
Protein post-translational modification identification. Panel A shows a bar plot of the ratio of confident PTMs identified in the study defined as the number of confident modification sites/number of total modification sites. Panel B illustrates the total number of PTMs in each technical replicate. Panel C shows the average expression of HPro on individual collagen chains between the three groups. In panel B, orange bars = N-acetyl-hexosamine (HexNAc), green bars = N-acetyl-hexosamine-hexosamine (HexNAcHex), blue bars = hydroxylysine (HLys) and purple bars = hydroxylproline (HPro). Red bars = DNH, green bars = DNS and blue bars = KTRH.

### Peptide level post-translational modification analysis elucidates collagen chains with highly abundant hydroxyproline and hydroxylysine modifications

Due to the abundance of HPro and HLys in collagens on the protein level, we wanted to investigate peptide-level confident HPro and HLys. Due to identical HPro and HLys mass shifts (+15.99), ambiguous PTM assignment is highly probable. This is more of an issue if a peptide contains multiple HPro and HLys sites that are near each other. Thus, we included HPro and HLys peptide chains with AScores>20, corresponding to a p-value of 0.01 (44). Furthermore, consistent with protein analysis, HPro and HLys peptides had to be abundant in 70% of samples in a biological group. In our pilot study, 54 HPro peptides and five HLys peptides met our criteria with COL1A2 and COL4A2 containing the most HPro and HLys peptides respectively. These peptides are listed in **Supplementary Excel File 6**. We calculated the weighted averages of confident HPro and HLys PTMs on collagen chains (**Figure 5**).

**Figure 5:**
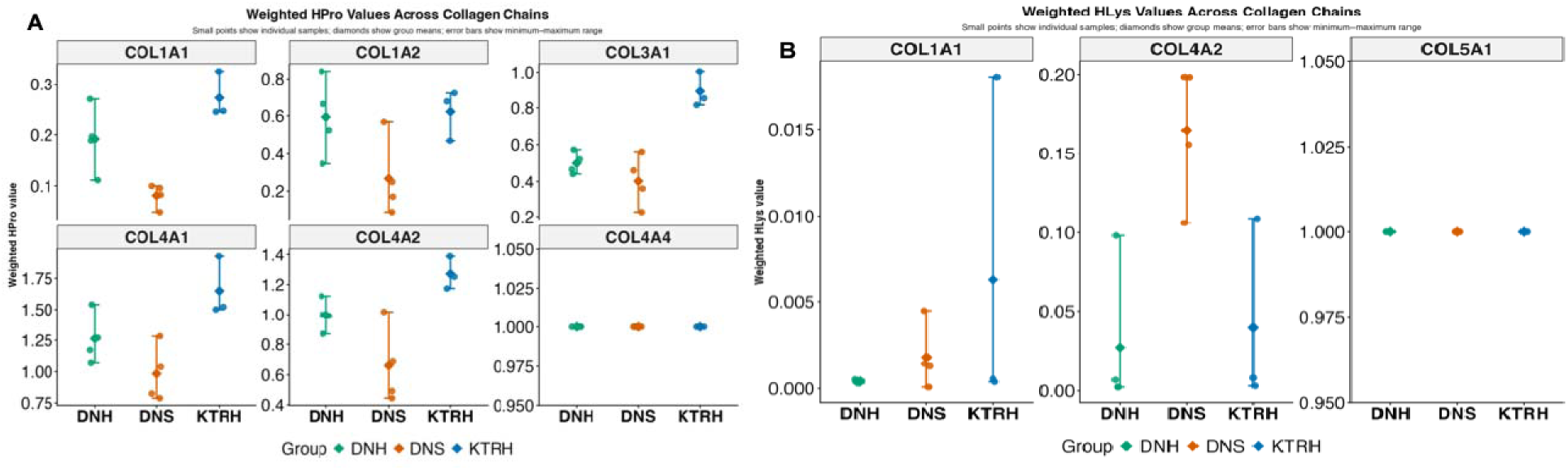
Abundances of HPro and HLys on collagen chains. Panel A illustrates the average weighted confident HPro values within each group across collagen chains. Panel B represents HLys average weighted values. Green diamonds indicate the DNH group, orange indicate DNS and blue KTRH. Error bars represent the range of weighted averages. In each panel, small points represent individual samples, diamonds are group means and error bars show the range (maximum-minimum).

The calculated weighted averages consider confidently modified (AScore>=20) and unmodified (AScore<=20 or unmodified) HPro and HLys and reflect the significance of an observation in each group. Our pilot results indicated that KTRH replicates had the highest HPro weighted averages amongst the three groups, thus these peptides possessed greater weights (or abundances) associated with HPro quantitation. Additionally, KTRH replicates had a stringent HPro range compared to DNS and DNH groups, except for COL1A1 and COL4A4, highlighting that most peptides possess consistent HPro modifications within collagen chains. Similar ranges between groups were observed for COL4A1. COL4A4 only had one peptide that contained a confident HPro modification and had uniform weighted averages across groups. Higher ranges/variability for DNS and DNH indicated HPro was inconsistent between collagen chains and DNH and DNS regions caused sporadic HPro. This finding could be necessary for regulating renal physiology in the presence of ECM dysregulation and disease pathogenesis. All HPro modifications were 4-hydroxyprolinated and not 3-hydroxyprolinated. Confident HLys sites (AScore>=20) were much fewer compared to HPro, and only a couple of COL1A1, COL4A2, COL5A1 chains generated them. The number of unmodified chains were also considered, yet due to the absence of HLys, their weights were non-existent. Peptide abundances associated with HLys were also less significant. KTRH reflected a higher range of HLys in COL1A1, while equal ranges were observed in COL4A2 and COL5A1. Similar to COL4A4, COL5A1 also possessed only one peptide that containing a confident HLys. Thus, a greater number of confidently modified HPro peptides contributed to the higher weights observed compared to HLys, indicating the significance of HPro in renal pathophysiological processes. However, due to the pilot nature of our study, it is difficult to discern whether these PTM alterations truly reflect a biological change due to disease, or if they occur by chance. Thus, it would be critical to determine the number of biospecimens required in each biological group to achieve sufficient power at a false discovery rate of 5%.

### Statistical analysis highlights compartmental-specific proteins driving separation of biological groups

Prior to running a robust power analysis, we first determined which proteins in each compartment contributed a large or small effect or magnitude when comparing DNS, DNH and KTRH groups. These effects would insinuate which proteins contribute the most (or least) to a group. **Table 2** illustrates proteins contributing the largest effect in each comparison (DNS vs DNH, DNH vs KTRH and DNS vs KTRH) in glomerular, tubulointerstitium, and “other” compartments respectively.

**Table 2:** Effect sizes and fold changes of compartmental-specific proteins. Top 5 proteins in each comparison (DNS-DNH, DNH-KTRH, DNS-KTRH) contributing a large positive Hedge’s g effect in the glomerulus, tubulointerstitium and “other” compartment. Values in bold are the average log2 fold change or Hedge’s g values for a compartment.

| Compartment | Average of Hedge’s g | Log2FC |
| --- | --- | --- |
| **<u>Glomerulus</u>** |  |  |
| **DNH-KTRH** | **1.75** | **4.43** |
| DYNC12 | 1.37 | 3.81 |
| KRT1 | 1.27 | 3.12 |
| PRDX2 | 1.52 | 1.54 |
| SERPINB6 | 3.72 | 10.42 |
| YWHAЕ | 0.85 | 3.28 |
| **DNS-DNH** | **2.15** | -5.70 |
| EZR | 4.37 | -7.20 |
| HSPA8 | 1.35 | -4.29 |
| SERPINB6 | 1.86 | -7.43 |
| TUBB2A | 1.58 | -4.80 |
| TUBB2B | 1.58 | -4.80 |
| **DNS-KTRH** | **3.50** | **6.83** |
| APCS | 3.72 | 8.31 |
| FBN1 | 5.08 | 6.51 |
| HBZ | 2.19 | 3.70 |
| LAMB2 | 3.82 | 8.98 |
| LGALS38P | 2.73 | 6.66 |
| **<u>Other</u>** |  |  |
| **DNH-KTRH** | **3.15** | **5.58** |
| TUBB4B | 5.13 | 9.27 |
| ATP5ME | 3.01 | 6.27 |
| ARPC4 | 2.93 | 8.04 |
| TALDO1 | 2.52 | .97 |
| RIDA | 2.16 | 3.37 |
| **DNS-DNH** | **7.79** | **-8.15** |
| NHERF1 | 9.61 | -9.06 |
| DHRS4L2 | 8.25 | -8.49 |
| HPD | 7.81 | -7.03 |
| ALDH7A1 | 6.84 | -8.23 |
| ACOT2 | 6.47 | -7.98 |
| **DNS-KTRH** | **6.08** | **10.44** |
| IGHA1 | 7.35 | 13.66 |
| DEUP1 | 6.64 | 10.16 |
| COL6A2 | 6.48 | 9.86 |
| ARPC4 | 5.02 | 8.94 |
| ADH1A | 4.94 | 9.62 |
| **<u>Tubulointerstitium</u>** |  |  |
| **DNH-KTRH** | **5.17** | **8.45** |
| FBP1 | 5.82 | 8.71 |
| AOC3 | 5.76 | 9.10 |
| PCBD1 | 5.22 | 9.04 |
| COL2A1 | 4.61 | 7.85 |
| GCDH | 4.48 | 7.76 |
| **DNS-DNH** | **11.25** | **-8.77** |
| ANPEP | 13.07 | -10.40 |
| SORD | 12.72 | -8.75 |
| CYB5A | 10.97 | -9.32 |
| PEPD | 10.42 | -6.78 |
| DDC | 9.91 | -8.61 |
| **DNS-KTRH** | **5.71** | **10.28** |
| PIGR | 7.63 | 11.67 |
| UMOD | 5.62 | 10.02 |
| SLC12A1 | 5.29 | 9.99 |
| COL2A1 | 5.07 | 10.18 |
| AOC3 | 4.94 | 9.56 |

An effect size > 0.2 was characterized as small, > 0.5 as moderate and > 0.8 as large, as described in the literature (56). In the glomerulus, EZR and FBN1 contributed the largest effects in the DNS-DNH and DNS-KTRH comparisons respectively, while SERPINB6 produced a large effect in DNH-KTRH comparison. UMOD and members of the solute carrier family (SLC12), are signature tubulointerstitium proteins and both exhibited large effects in the DNS-KTRH comparison. In the DNH-KTRH comparison, AOC3 and FBP1 resulted in similar large effects as the DNS-KTRH comparison. However, larger effects were observed amongst all proteins in the DNS-DNH comparison (ANPEP, SORD, CYB5A), signifying the importance of these proteins in driving separation between comparisons. In the “other” category, effects were largest in the DNS-DNH comparison similar to the tubulointerstitium, **Supplementary Excel File 7** contains the list of protein Hedge g’s effect sizes in each comparison. Using the Hedge’s g effect sizes and log2 fold changes, we created our limma model for power analysis. the number of samples, based on the effect sizes, required to observe a statistical significance difference of FDR < 0.05 and a power level > 70%. **Figure 6** illustrates comparison of power (**Figure 6A**) and FDR (**Figure 6B**) across comparisons.

**Figure 6:**
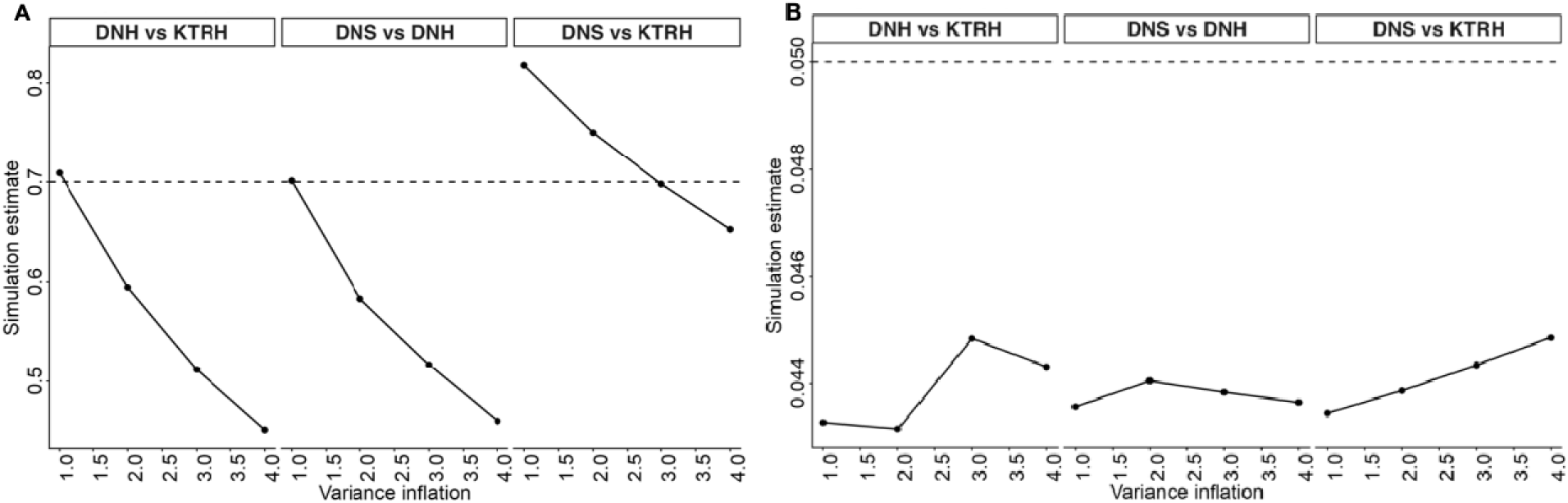
Power analysis. Power level fluctuations are represented in each comparison in panel A, while FDR values are illustrated in panel B. Variance factors are plotted along the x-axis, and each individual dot represents a variance inflation value. The dashed lines in panels A and B represent the target power level and FDR respectively.

After running our limma simulation over 1000 iterations, we determined that ∼51 independent samples are required in all three comparisons to achieve the desired FDR and power levels. As shown, DNS-KTRH comparison had the achieved the highest power (80%) even with variance inflation factors at 2 and 3, highlighting that potential intra-variability is lower compared to feature abundances in the DNH-KTRH and DNS-DNH comparisons. DNH-KTRH and DNS-DNH comparisons exhibited a rapid decrease of power as variance inflation increased, thus requiring a higher number of samples to achieve the target values. FDR levels between comparisons were lowest at a variance inflation factor of 1; however, only DNS-KTRH showed a linear increase in FDR as variance inflation increased, while DNH-KTRH and DNH-KTRH showed non-linear patterns. Follow-up studies would need to be performed with the suggestive larger sample size to observe whether changes are statistically significant and biologically meaningful.

## Discussion

We illustrated unique proteomic signatures within each group and pinpointed compartmental-specific proteins (e.g. tubulointerstitium and glomerulus). driving separation in each comparison. The term “tubulointerstitium” is used ubiquitously in the kidney literature to describe the shared tubule and interstitium microenvironment, and there could be potential proteins that are significantly enriched in the tubules and secreted into the interstitium during renal (patho)physiology compared to those proteins that are strictly localized to the tubules or interstitium. Due to the paucity of proteomic (or omic) studies investigating kidney vasculature, we did not include it in our study. Proteins that were not significantly enriched (or upregulated) in one compartment nor categorized as signature proteins were categorized as “other”.

We analyzed peptide level expression amongst the top peptides across replicates and quantified PTM-specific patterns on matrisome proteins in each group, highlighting unique PTM abundances contributing to renal pathophysiology. Due to the exploratory nature of this study and low sample size in DN and KTRH groups, we generated a model to predict the required number of samples necessary to obtain 70% power and a FDR < 0.05 per group. We also evaluated the reproducibility of proteomics signatures in the tubulointerstitium from replicates of healthy and chronically damaged tubulointerstitium. Quality control analyses such as PCA and correlation highlight intra-group homogeneity in protein expression, demonstrating reproducibility in our LMD experimentation.

We quantified signature compartment proteins in the tubulointerstitium across DNS, DNH and KTRH tubulointerstitium. Most of the quantified proteins were either enriched in a compartment yet were not signature- or compartment-specific, highlighting their diverse roles in organ physiology. Several proteins that were enriched in the glomerulus did not exhibit significant alterations between groups such as KRT10, COL4A4, PRXD2. Under baseline healthy conditions, KTR10 is not expressed, yet it is a regulator of glomerular crosstalk during inflammatory diseases (e.g., IgA nephropathy) (57). COL4A4 abundance is a major network-forming collagen responsible for providing structural and tensile support; however, it was particularly enriched in the GBM, though it is also found in the tubules as well (26). Collagen type IV has been extensively studied in diabetic nephropathy and is associated with renal function and glomerular pathologies (58, 59). Other proteins such as laminin (LAMB2), FBN1 and APCS were highly abundant in the DNS samples. LAMB2 binds to cell surface receptors such as integrins and mediates cell-ECM interaction and is a major component in the mature GBM (60). FBN1 induces mesangial expansion promoting glomerulosclerosis (61), and APCS is a component of the glomerular basement membrane. Interstitial-enriched proteins were sparse in our data, thus only a subset of interstitial proteins were quantified. HSPG2, a heparan sulfate proteoglycan, was abundant across all DNS and KTRH replicates. Heparan sulfate proteoglycans have been studied in the tubulointerstitium and regulate immunity (62), but they are also a major component of the GBM (63). Tenascin-C was more abundant across DNS replicates compared to KTRH and DNH replicates. Tenascin-C, a matrisome glycoprotein, impacts numerous cellular mediators such as integrin αvβ6, EGFR, Wnt and TLR4 promoting immune recruitment and fibrosis in the kidney (64). It has been shown to localize to the tubulointerstitium and glomerulus during CKD and is upregulated in DN studies (65, 66). Regarding classical tubular signature proteins, CUBN (67), LRP2 (67), SLC family members (SLC6A19, SLC3A2, SLC4A4) (68) and PDZK1 were quantified in our data. PDZK1 has been shown to be a mitigator of fibrosis (69). Cubulin and megalin are two proximal tubulin receptors responsible for albumin absorption, and dysfunction of these receptors during DN could lead to microalbuminuria (70). Most of these proteins were more abundant in healthy tubulointerstitium (DNH, KTRH) compared to chronically damaged tubulointerstitium, demonstrating their role in regulating ion and amino acid transport and internalization during renal physiology. Their decrease abundance in chronically damaged tubulointerstitium could indicate the initiation of interstitial matrix, presence of immune cells in the interstitial matrix and tubular atrophy, thus deteriorating core tubular transport functions. required in the kidney. It is important to note that proteins categorized as “other” in our does not indicate their complete absence in a kidney compartment. These proteins could have low abundance and thus be below our instrument’s limit of detection. Overall, the tubulointerstitium compartment contained the most signature marker proteins and a higher number of enriched proteins compared to other compartments as expected. Thus, our LMD methodology consisted of pure replicates of a compartments, yet some glomerular material might have been incidentally captured. Section thickness is non-zero, and a small percentage of background cells (e.g., glomerulus) could lie beneath the tubulointerstitium.

Traditional proteomics experiments involve inferential statistics to determine differentially expressed proteins (DEPs) between biological groups. Studies typically use a fold change and adjusted FDR p-value cutoff to identify DEPs. However, emerging studies are suggesting that researchers report other statistical metrics (confidence intervals, effect sizes) to determine the differences in mean between two groups and evaluate the significance of a study and (49). R packages are beginning to incorporate the Cohen d effect sizes to help scientists systematically determine DEPs in their studies (50). Effect sizes have been reported for studies in diabetic retinopathy (71), kidney transplant patients with COVID-19 (72) and neural stem cells (73). Since our study had only N=1 per group, inferential statistics would violate underlying assumptions regarding the data. Thus, we incorporated effect sizes to determine which proteins could contribute to driving biological group separation, and future large-scale studies would be able to determine statistical significance. As shown, effect sizes between comparisons varied widely. Glomerular effect sizes ranged from 0.85-5.08, while tubulointerstitium effect sizes were from 4.48-13.07. These effect sizes correlated with the log2 fold changes for these proteins; tubulointerstitium proteins had a higher average log2 fold change in each biological group compared to glomeruli. In both compartments, the DNS-KTRH comparison exhibited the highest average log2 fold change and average Hedge’s g effect size, indicating that these proteins potentially contribute greatly to the morphological and histological changes observed in tubulointerstitium between DNS and KTRH patients, warranting further investigation. However, it is important to note that these effect sizes (and log2 fold changes) are overestimates since they were sampled from technical and not biological replicates. Variance between technical replicates could be artificially low and less stable, and thus falsely inflate effect sizes and fold changes to reflect significance. Additionally, we determined that a minimum of 50 samples are required to achieve a power of 70% and a FDR < 0.05.

Hydroxylproline was identified as the most prevalent PTM on the protein and peptide level, specifically on collagens, and 4-Hydroxyproline was the predominant pattern identified across collagen chains, corroborating literature findings (74–76). We found a greater, weighted abundance of confident hydroxyproline quantitation across collagens (CO1A1, COL3A1, COL4A1, COL4A2) in KTRH replicates compared to DNS and DNH replicates. Kidney transplant patients are a unique subset of CKD patients. They are typically classified as CKD due to their suboptimal eGFR (<60 min/mL/1.73 m^2^), potential high proteinuria and overall risk of kidney failure if the transplant is not closely monitored. Additionally, the probability of developing fibrosis increases from 25% at one-year post-transplant to 65% at 2-5 years, thus fibrosis correlates with decreased transplant survival (8). One recent study assessed hydroxyproline content in urine microvesicles in kidney transplant recipients with fibrosis (40). They observed an increase in urinary hydroxyproline in kidney transplant patients with fibrosis compared to healthy patients and noticed an even greater amount in patients with inflammation, thus identifying hydroxyproline as a potential biomarker of fibrosis. Thus, our findings of increased hydroxyproline abundance in healthy tubulointerstitium in kidney transplant patients compared to other groups support this prior study and could indicate hydroxyproline as a novel biomarker in these patients as well. To our knowledge, there have not been additional studies investigating hydroxyproline as a biomarker of fibrosis in kidney transplant patients or diabetic nephropathy patients.

A major limitation of this study was the small sample size and the lack of age-sex match samples per group. Since this study had N=1 for two different biological groups it is impossible to perform differential expression analysis and evaluate true biological variation in protein expression between groups and deduce strong conclusions. The KTRH biopsy was from a 47-year-old male patient, while the DN case came from an 86-year-old female. Aging can lead to reduction in kidney cortical volume, increase in medullary volume and interstitial fibrosis and tubular atrophy (77). Thus, though we are certain that IFTA was caused by disease, especially in the older DN patient exhibiting advanced chronic changes, we are doubtful about whether the IFTA was a direct result of DN or aging during the collection of chronically damaged tubulointerstitium. Additionally, it is difficult to determine whether PTM alterations reflect true variation due to disease or by chance without sufficient power and deep experimental investigation of any changes in a PTM’s site localization and abundance during disease. We also did not include a control group in this study to compare baseline expression of compartmental-specific proteins with those in the diseased (KTRH and DN) groups. A suitable control group would be native kidney biopsies with minimal IFTA and the absence of any critical pathologies that would confound our results. Furthermore, we did not LMD KTRH chronically damaged tissue which would enhance our experimental design by enabling the comparison of KTRH groups and discovering changes in tubule-interstitial matrisome signatures characterizing fibrosis. We chose DN since our BMC repository contains more DN cases than any other kidney disease, and it is a ubiquitously studied renal model. KTRH samples were more difficult to procure, especially those that have not undergone significant fibrosis nor have any other underlying condition. Future studies should also incorporate high power, in-depth PTM peptide analysis on matrisome and intracellular proteins in common kidney diseases and investigate how these PTMs are significantly changing between biological phenotypes in early- and late fibrosis. Results from these studies would lead to the elucidation of novel molecular mechanisms that contribute to IFTA and discovery of new targets that could mitigate and reverse fibrosis in kidney transplant patients and non-transplant CKD patients.

## Data Availability

The mass spectrometry proteomics data have been deposited to the ProteomeXchange Consortium via the PRIDE (78) partner repository with the dataset identifier PXD079025 and 10.6019/PXD079025.

## Declaration of generative AI and AI-assisted technologies in the manuscript preparation process

During the preparation of this work, the author used ChatGPT to aid in the creation of limma model to simulate the number of biospecimens required to obtain an FDR < 0.05 and power of 70% for a future study. After using this tool/service, the author(s) reviewed and edited the content as needed and take(s) full responsibility for the content of the published article. The code is provided in **Supplemental File 1**.

## Author Contributions

J.C.L. and J.Z. conceptualization; J.C.L. data curation; J.C.L. A.C. and M.S. formal analysis; J.C.L. writing-original draft; J.C.L., J.H., J.Z. writing-reviewing and editing; J.C.L., J.H., J.Z. investigation; J.C.L, J.H., J.Z. methodology; J.Z. project administration; J.C.L. software; J.C.L. visualization; J.H. and J.Z. resources; J.Z. funding acquisition; J.H. and J.Z. supervision.

## Acknowledgements

This work was supported by NIH grants R01AG075876 and R35GM144090.

